# UCHL1-dependent control of Hypoxia-Inducible Factor Transcriptional Activity in Liver Disease

**DOI:** 10.1101/2023.01.08.523142

**Authors:** Amy Collins, Rebecca Scott, Caroline L Wilson, Giuseppe Abbate, Gabrielle Ecclestone, Demi Biddles, Fiona Oakley, Jelena Mann, Derek A Mann, Niall S. Kenneth

## Abstract

Liver fibrosis is the excessive accumulation of extracellular matrix proteins that occurs in most types of chronic liver diseases. Fibrosis is associated with the activation of hepatic stellate cells (HSCs) which transdifferentiate into a myofibroblast like phenotype that is contractile, proliferative and profibrogenic. Hypoxia-inducible factor 1 (HIF1), an oxygen-sensitive transcription factor, is elevated during HSC activation and promotes the expression of profibrotic mediator HIF target genes. HIF activation during HSC activation can by either due to localised decreases in oxygen levels, or through oxygen-independent mechanisms that are not completely understood. Here we describe a role for the deubiquitinase UCHL1 in regulating HIF levels and activity during HSC activation and liver fibrosis. Increased HIF1α expression correlated with induction of UCHL1 mRNA and protein with HSC activation. Genetic deletion or chemical inhibition of UCHL1 impaired HIF activity through reduction of HIF1α levels. UCHL1 specifically cleaves the degradative ubiquitin chains from HIF1α leading to increased HIF1α levels, even in sufficiently oxygenated cells. Furthermore, our mechanistic studies have shown that UCHL1 elevates HIF activity through specific cleavage of degradative ubiquitin chains, elevates levels of pro-fibrotic gene expression and increases proliferation rates. These results demonstrate how small molecule inhibitors of DUBs can modulate the activity of HIF transcription factors in liver disease. Furthermore, inhibition of HIF activity via modulation of the ubiquitin-proteasomal degradation pathway may represent a therapeutic opportunity with other HIF-related pathologies.

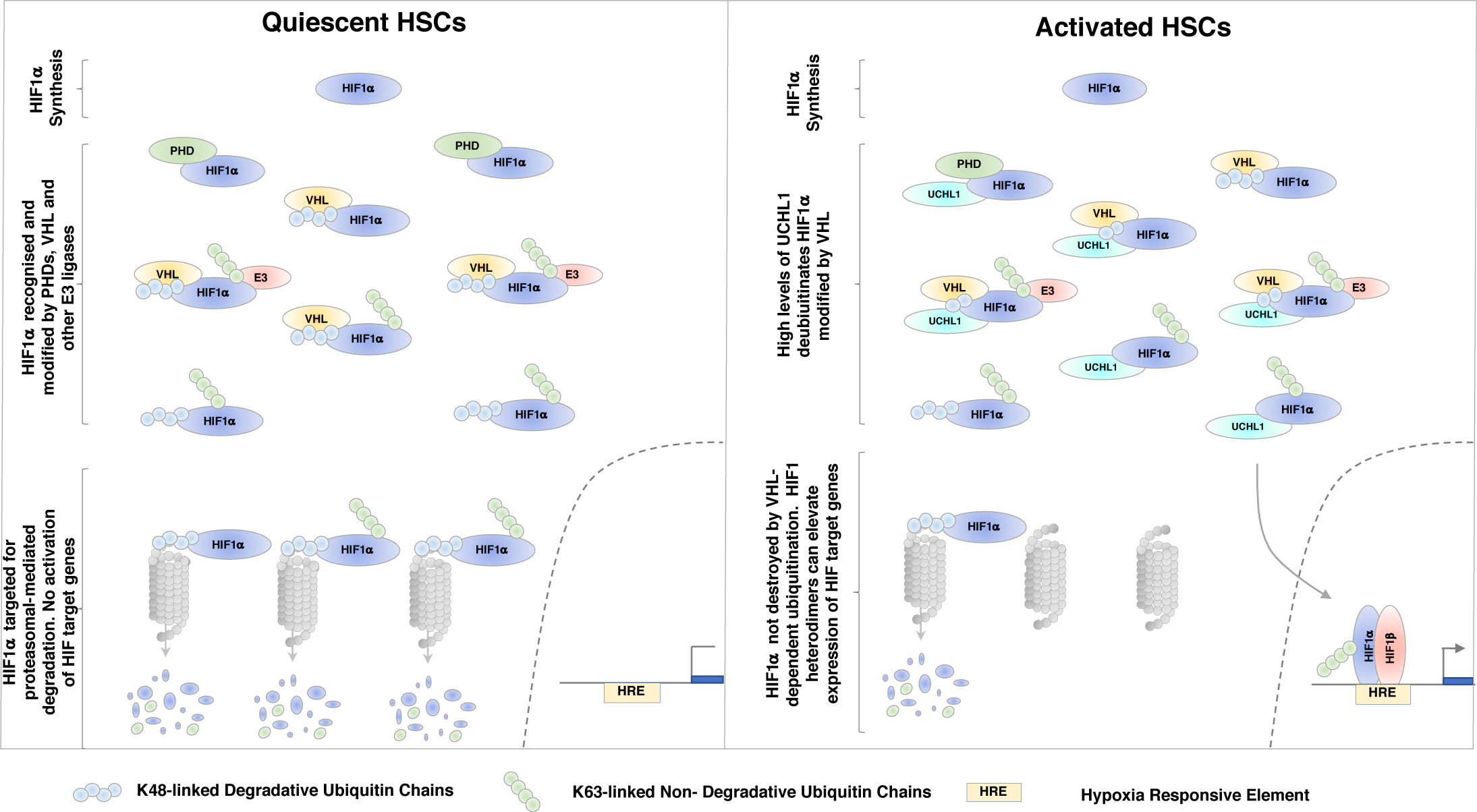

## Introduction

Hypoxia Inducible Factor 1 (HIF1) is a ubiquitously expressed heterodimeric transcription factor composed of an oxygen-sensitive alpha subunit and a constitutively expressed beta subunit, also called aryl hydrocarbon receptor nuclear translocator (HIF1*β*/ ARNT) [1, 2]. In sufficiently oxygenated cells HIF1α is rapidly hydroxylated by a family of prolyl-hydroxylases (PHD1, PHD2, or PHD3) [1, 2]. Hydroxylated HIF1α is then recognized by a multimeric protein complex that includes the product of the Von Hippel Lindau (VHL) tumour suppressor gene, an E3 ubiquitin ligase [2, 3]. Prolyl hydroxylation and subsequent VHL-dependent ubiquitination leads to rapid proteasomal degradation and clearance of HIF1α subunits [3]. In normal oxygen conditions the half-life of HIF1*α* is <5min [3]. When oxygen is limiting PHDs are inhibited and HIFs escape hydroxylation and dimerise with ARNT [3]. Active HIF heterodimers can then bind to hypoxia-responsive elements (HREs) in the promoter region of target genes to activate transcription [3]. HIF1 promotes the expression of a broad range of target genes including the expression of genes that increase oxygen delivery, which enables cells to survive in oxygen-deficient conditions [3].

Clinical and experimental data suggest that HIF activity is disrupted in multiple human pathologies such as heart disease, cancer, cardio vascular disease and chronic obstructive pulmonary disease [4]. Activation of HIF and hypoxia-dependent signalling cascades is also emerging as a regulator of liver fibrosis [5]. Hepatocyte-specific deletion of HIF1α results in significant decreases of pro-fibrotic mediator levels, such as PDGFs, FGFs and growth factors leading to reduced ECM deposition and a reduction of the fibrotic phenotype [6]. Conversely, targeted genetic deletion of negative HIF regulators such as PHDs and VHL results in high HIF levels and increase hepatic steatosis and fibrosis [7, 8]. These genetic studies implicate a role for HIF signalling in the biology of the hepatic stellate cell (HSC). The transdifferentiation of quiescent HSC (qHSC) to activated (aHSC) myofiroblasts is pivotal in liver fibrosis, with the aHSC being responsible for deposition of collagen-rich fibrotic matrix [9, 10]. Proliferation and migration of aHSCs is important in the progression and spread of fibrosis in the diseased liver. The precise mechanism(s) by which HIF influences HSCs and fibrogenesis is unclear [5]. Regions of hypoxia can develop in the liver after acute liver injury, however, increased HIF1α levels and activity are observed in diseased liver before the development of hypoxia, suggesting an oxygen-independent mode of HIF activation in chronic liver disease [11, 12].

HIF activity can be elevated through the up regulation of factors that disrupt HIF1α degradation, even in the presence of sufficient oxygen [13, 14]. Deubiquitinating enzymes such as the Ubiquitin C-terminal hydrolase-L1 (UCHL1) promote HIF1 activity under normoxic and hypoxic conditions by disrupting ubiquitin-mediated, proteasomal degradation of HIF1α [15]. Increased expression of UCHL1 is associated with elevated HIF1α levels and distant metastasis in cancer patients, suggesting that UCHL1 may promote metastasis through elevation of HIF1 activity [15–17]. Interestingly, analysis of DUB expression during HSC activation reveals that UCHL1 is highly induced during HSC activation, perhaps indicating a mechanism by which HIF1 activity is elevated during liver fibrosis [18].

In this present study, we examined the role of UCHL1 in regulating HIF1 activity during HSC activation, and the role of UCHL1 in driving the expression of hypoxia- and pro-fibrotic gene expression in a tractable cellular model. HIF1α levels were elevated in aHSC. Furthermore, genetic deletion or chemical inhibition of UCHL1 markedly reduced HIF1α levels under both normoxic and hypoxic conditions. Expression of UCHL1 elevated levels of hypoxia-responsive and pro-proliferative/fibrogenic gene expression in a HIF-dependent manner by specifically deubiquitinating degradative ubiquitination from the HIF1α subunit. As an inhibitor of UCHL1 was able to suppress fibrogenesis in a human precision cut liver slice (hPCLS) model of fibrosis targeting UCHL1/HIF may be an interesting strategy to explore towards the prevention of fibrosis progression in chronic liver disease.

## Materials and Methods

### Isolation of primary HSC

Primary human HSCs were isolated from normal margins of surgically resected liver. Liver tissue was digested with pronase and collagenase B (Roche) and the cell suspension was subsequently separated by an 11.5% Optiprep gradient (Sigma). HSCs were seeded onto plastic (Corning), cultured in Dulbecco’s modified Eagle’s medium (Life Technologies) supplemented with 16% fetal bovine serum, pyruvate, glutamine, penicillin, and streptomycin (Life Technologies) and maintained in an incubator at 37 °C with 5% CO_2_. Freshly isolated (day 0) cells were considered quiescent and (day 10) cultures regarded as activated.

Mouse HSC were isolated from normal livers of *UCHL1*^-/-^ or WT littermate controls. Liver tissue was digested with pronase and collagenase B (Roche) and the cell suspension was subsequently separated by an 11.5% Optiprep gradient (Sigma). HSCs were seeded onto plastic (Corning), cultured in Dulbecco’s modified Eagle’s medium (Life Technologies) supplemented with 16% fetal bovine serum, pyruvate, glutamine, penicillin, and streptomycin (Life Technologies) and maintained in an incubator at 37 °C with 5% CO_2_.

HEK293 cells were maintained in DMEM, supplemented with 10% FBS, glutamine, penicillin, and streptomycin (Life Technologies) and maintained in an incubator at 37 °C with 5% CO_2_. HEK293-HRE-luciferase cells were a kind gift from Professor Sonia Rocha, Liverpool, and maintained in DMEM, supplemented with 10% FBS, 0.5 µg/ml puromycin, glutamine, penicillin, and streptomycin (Life Technologies) and maintained in an incubator at 37 °C with 5% CO_2_.

### DNA constructs and transfections

pcDNA3-HA-HIF1α (Addgene #18949), pcDNA3-HA-HIF1α P402A, P564A (Addgene #18955) were gifts from William Kaelin supplied by Addgene. pEBB. pEBB Flag VHL, pEBB-N-biotin-HIF1α and pEBB-His-ubiquitin have been previously described[19]. Flag-HA-UCHL1 (Addgene # 22563) was a gift from Wade Harper, supplied by Addgene. pEBB-HA-UCHL1 was subcloned from Flag-HA-UCHL1 into pEBB-HA. HEK293 transfections were performed using a standard calcium phosphate method [19].

### Cell lysis and Immunoblotting

Cells were lysed in RIPA lysis buffer and immunoblotted as described[20]. Antibodies used were human HIF1α (Clone 241809, R&D systems), rodent HIF1a (#14179, Cell Signaling Technologies), Hydroxy-HIF-1α (Pro564) (#3434, Cell Signaling Technologies), VHL (#68547, Cell Signaling Technologies), ubiquitin (sc-8017, Santa Cruz), β-actin (AC-74, Sigma), HA (Clone 16B12, Covance), Flag (M2, Sigma), Biotin (sc-53179, Santa Cruz), UCHL1 (#3524, Cell Signalling Technologies).

### Ubiquitination assays

For ubiquitination assays, cells were lysed at room temperature under denaturing conditions (8 M urea, 50 mM Tris [pH 8.0], 300 mM NaCl, 50 mM Na2HPO4, 0.5% NP-40, 1 mM PMSF, supplemented with protease inhibitors) and ubiquitinated material was recovered by rotation with NiNTA-agarose (Invitrogen) washed 3× with lysis buffer and analysed by western blotting [19].

### Co-precipitation Assays

48 hours following transfection cells were harvested in cell lysis buffer (20mM Tris pH 8.0, 150mM NaCl, 1mM EDTA, 1% Triton X-100, supplemented with protease and phosphatase inhibitors). Biotinylated HIF1α was purified with streptavidin agarose beads (Sigma) and precipitates analysed by immunoblot analysis[19].

### Quantitative reverse transcription-PCR

Total RNA was isolated using the Peqgold Total RNA Isolation Kit (Peqlab) according to the manufacturer’s instructions. Reverse transcription with random and oligo(dT) primers and MMLV reverse transcriptase (Quanta Biosciences) was performed on 500 ng of total RNA.

Quantitative PCR data was generated using the following experimental settings: hold 50°C for 3 min; Hold 95°C 10 min; cycling (95°C for 30 s; 58°C for 30 s; 72°C for 30 s with fluorescence measurement for 45 cycles). All values were calculated relative to maximum hypoxic induction and normalised to RPL13A levels using the Pfaffl method. Each cDNA sample was assayed in duplicate and the results shown are averages derived from three or four biological repeats with error bars indicating the standard deviation.

Primers for human samples were: RPL13A sense 5′-CCT GGA GGA GAA GAG GAA AGA GA-3′, antisense 5′-TTG AGG ACC TCT GTG TAT TTG TCA A-3′; ANKRD37 sense 5′-GTC GCC TGT CCA CTT AGC C-3′, antisense 5′-GCT GTT TGC CCG TTC TTA TTA CA-3′; CAIX sense 5′-CTT TGC CAG AGT TGA CGA GG-3′, antisense 5′-CAG CAA CTG CTC ATA GGC AC-3′; COLA1 sense 5’-ATG TGC CAC TCT GAC TGG AA – 3’, ANTISENSE 5’-CTT GTC CTT GGG GTT CTT GC-3’; GLUT1 sense 5′-CTG GCA TCA ACG CTG TCT TC-3′, antisense 5′-GCC TAT GAG GTG CAG GGT C-3′; HIF1α sense 5′-CAT AAA GTC TGC AAC ATG GAA GGT-3′, antisense 5′-ATT TGA TGG GTG AGG AAT GGG TT-3′; SMA1 sense 5’-ACC CAG CAC CAT GAA GAT CA-3’, antisense 5’-TTT GCG GTG GAC AAT GGA AG -3’; TGF*β* sense 5’-CGT GCT AAT GGT GGA AAC CC-3’, antisense 5’-TCG GAG CTC TGA TGT GTT GA-3’; TIMP1 sense 5’-GTT TTG TGG CTC CCT GGA AC-3’, antisense 5’-GTC CGT CCA CAA GCA ATG AG-3’; VEGF sense 5′-CCT GGT GGA CAT CTT CCA GGA GTA CC-3′, antisense 5′-GAA GCT CAT CTC TCC TAT GTG CTG GC-3′.

Primers for murine samples were Hif1a sense 5’-CCT GCA CTG AAT CAA GAG GTT GC-3’, antisense 5’-CCA TCA GAA GGA CTT GCT GGC T-3’

### Proliferation experiment

Clonal cell lines were seeded into 96 well plates at a density of 2500 cells/well. The plates were incubated at 37°C and measured using 5% PrestoBlue® after either 24, 48, 72 or 96 hours’ incubation. The gain was determined after 24 hours and was subsequently used to measure all plates to allow for comparison between time points. Raw values were plotted to indicate cell growth/ proliferation over time. The data were plotted using GraphPad PRISM and the area underneath the curve was quantified. Statistical analysis was performed by Student’s T-test. A value of p<0.05 was considered statistically significant.

### Luciferase assays

Lysates for luciferase assay were prepared in 1× passive lysis buffer (Promega), 100 μl per well of a 24-well plate. 10 μl of lysate was incubated with 50 μl luciferase reagent (Promega) and measured for 10 s using (Lumat LB9507, EG&G Berthold).

Graphs represent raw RLUs readings from four independent experiments. Raw RLU values are presented, and statistical analysis performed using a one-way ANOVA with Dunnett’s multiple comparisons test. A value of p<0.05 was considered statistically significant.

#### Precision Cut Liver Slices

Human liver tissue was cored using a 8-mm Stiefel biopsy punch (Medisave, Weymouth, UK) and cut into 250micron slices using a Leica VT1200S vibrating blade microtome (Leica Biosystems, Milton Keynes, UK) as previously described [21]. All PCLSs were cultured in Williams medium E (W4128; Sigma-Aldrich), supplemented with 1% penicillin/streptomycin and l-glutamine, 1× insulin transferrin-selenium X, and 2% fetal bovine serum (Thermo Fisher Scientific, Cramlington, UK), and 100 nM of dexamethasome (Cerilliant, Texas, USA) at 37°C, supplemented with 5% CO2. Media were changed daily.

### Statistical analysis

Statistical analysis was performed by a one-way ANOVA with Tukey’s multiple comparisons test or unpaired Student’s t-test where appropriate. If a p-value is less than 0.05, it is flagged with one star (*). If a p-value is less than 0.01, it is flagged with two stars (**). If a p-value is less than 0.001, it is flagged with three stars (***).

## Results

### UCHL1 Deubiquitinates Degradative Ubiquitin Chains from HIF1α

HIF1 activity is mainly controlled by the availability of the oxygen labile HIF1α subunit. When sufficient oxygen is available HIF1α is destroyed by ubiquitin-mediated proteasomal degradation, however during hypoxia ubiquitination is prevented, allowing HIF1α to accumulate, bind with HIF1β, and form the active HIF1 complex [3]. However, HIF1α ubiquitination is reversible and ubiquitin chains can be removed through the action of deubiquitinating enzymes (DUBs) to exert fine control HIF1α stability [22].

Ubiquitin C-terminal hydrolase-L1 (UCHL1), is a DUB whose expression is associated with elevated HIF activity[15–17]. To begin to determine mechanistic relationships between UCHL1 and HIF, cells were co-transfection with tagged UCHL1, HIF1α and Ubiqutin (Ub) expression vectors (Figure 1). Despite the active site for UCHL1 appearing to limit access for proteins other than very short, ubiquitin-conjugated peptides, ubiquitination of HIF1α is markedly reduced when UCHL1 is co-expressed [23] (Figure 1A). Interestingly, expression of UCHL1 also resulted in deubiquitination of HIF2α, indicating that UCHL1 can affect both HIFα isoforms targeted by VHL-dependent ubiquitination, thus exerting control of the 2 major HIFα subunits (Figure 1B).

**Figure 1.**
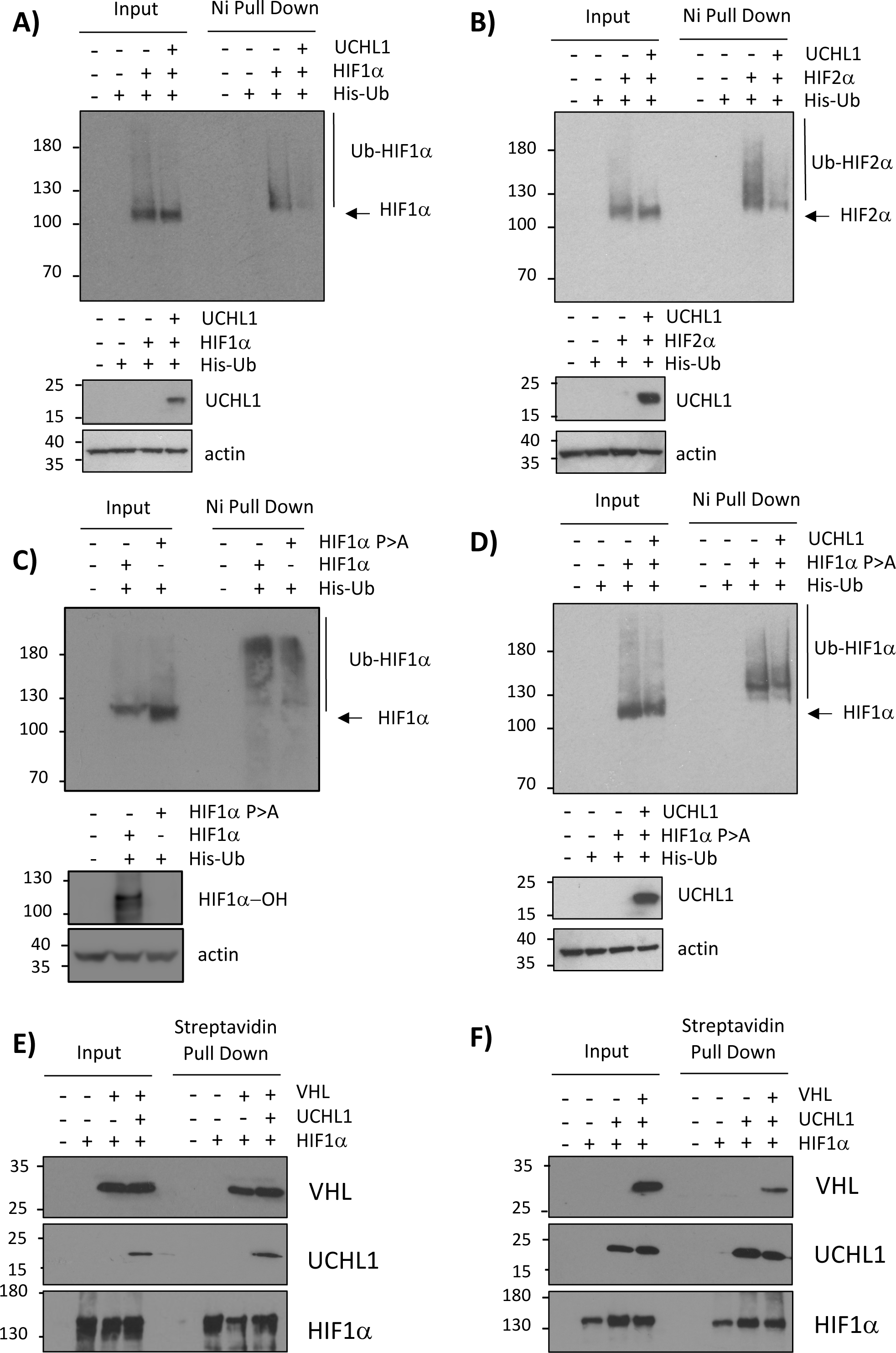
UCHL1 specifically degrades VHL-dependent Ubiquitin chains from HIF1α. **(A)** HEK293 cells were transiently transfected with plasmids encoding HA-UCHL1, His tagged ubiquitin along with HA-HIF1α. 48 hours post-transfection ubiquitinated complexes were stabilized by the addition of MG132 5h prior to harvesting. Ubiquitinated material was recovered from lysates by incubation with nickel-coated beads and analysed by immunoblotting. UCHL1 expression was confirmed by immunoblotting input lysates with a anti UCHL1 antibody. **(B)** As in (A) using HA-HIF2α **(C)** HEK293 cells were transiently transfected with plasmids encoding HA-HIF1α, HA-HIF1α (P402A, P562A) with His-tagged ubiquitin. Ubiquitin pull downs performed as in (A). **(D)** As in (A) using HA-HIF1α (P402A, P562A) **(E-F)** HEK293 cells were transiently transfected with plasmids encoding (N-terminal Biotin Tagged) NBT-HIF1α, HA-UCHL1 and Flag-VHL as indicated. 24 hours prior to harvesting growth media was supplemented with biotin (4uM). HIF1α precipitates were recovered from lysates by incubation with streptavidin coated beads and analysed by anti HIF1α, anti-Flag, anti-HA and anti-actin immunoblotting.

HIF1α is a highly ubiquitinated protein with over 20 known sites of ubiquitination (https://www.phosphosite.org/proteinAction.action?id=4987&showAllSites=true). In addition to being regulated by ubiquitin-mediated proteasomal destruction, HIF1α can be modified by non-degrative ubiquitin chains that can enhance its activity by modulating its levels and/or subcellular localization [22]. Ubiquitination assays reveal that a HIF1α mutant that is resistant to VHL-dependent ubiquitination (HIF1α P402A, P564A) is still ubiquitinated in cells at lower levels (Figure 1C). However, overexpression of UCHL1 does not reduce levels of HIF1α P402/P564A ubiquitination, suggesting that UCHL1 only prevents the accumulation of degradation-inducing ubiquitin chains onto HIFα subunits (Figure 1D).

### UCHL1 and VHL directly bind to HIF1α through independent mechanisms

As analysis of UCHL1 structure suggests that it is unlikely to directly cleave ubiquitin chains from substrates it has been proposed that UCHL1 decreases HIF1α ubiquitination by preventing the HIF1α / VHL interaction [15]. Co-precipitation experiments confirm that HIF1α precipitated efficiently with both VHL and UCHL1 (Figure 1E and 1F). However, neither UCHL1 or VHL could displace the other from HIF1α, indicating that they can bind to HIF1α simultaneously through distinct interaction domains, and UCHL1-depdendent deubiquitination of HIF1α is not achieved through prevention of the VHL/ HIF1α interaction (Figure 1E and 1F).

### Elevated UCHL1 enhances HIF activity

Having established that UCHL1 can reduce HIF1α ubiquitination we next investigated the relationship between UCHL1, HIF and hypoxia on HIF target gene expression. UCHL1 was expressed in HEK293 cells (a cell line that expresses no detectable levels of endogenous UCHL1) and exposed to hypoxia. Expression of UCHL1 increased the levels of hypoxia-induced HIF1α as compared to control cells as measured by immunoblot analysis (Figure 2A). HIF activity was assessed using HEK293 cells containing an integrated luciferase reporter construct possessing three copies of the hypoxia-responsive element (HRE) consensus-binding site. UCHL1 expression stimulated higher levels of luciferase activity under conditions of normoxia (Figure 2B), and hypoxia (Figure 2C) as compared to control cells, consistent with the UCHL1-dependent increase in HIF1α levels.

**Figure 2.**
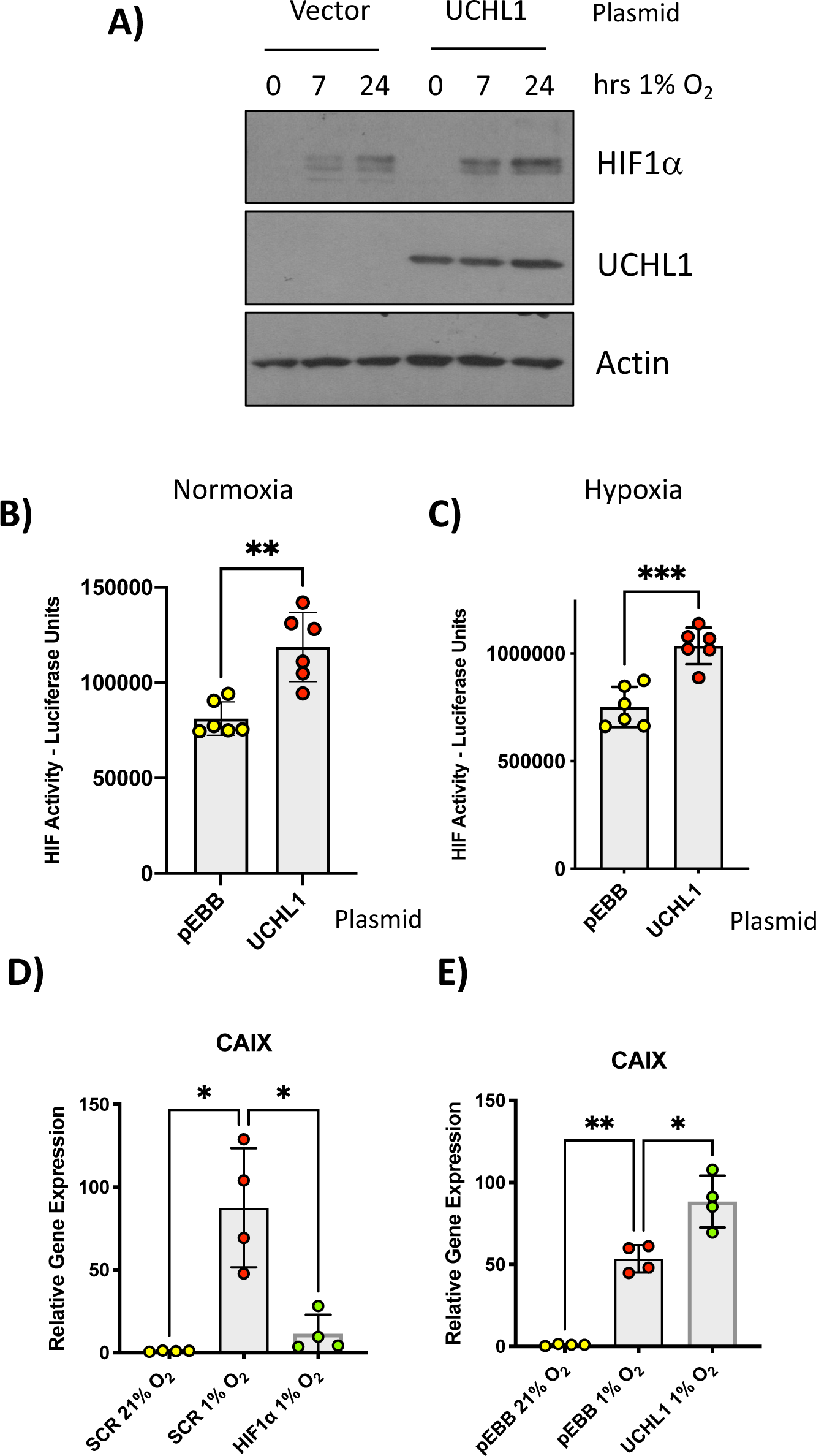
UCHL1 activates HIF-dependent gene expression. **(A)** HEK293 cells transfected with the indicated plasmids before being exposed to 1% O2 for the indicated times. Whole-cell lysates (WCLs) prepared from these cells were subjected to immunoblot analysis to assess expression levels of the indicated proteins. **(B)** HEK293 cells stably expressing luciferase driven from the HRE promoter element (HRE-Luc) were transfected with UCHL1 and luciferase activity determined 48 hours post-transfection. **(C)** as in (B) with cells incubated in 1% O_2_ 24h prior to lysis. **(D)** Quantitative RT–PCR analysis of CAIX mRNA prepared from HEK293 cells transfected with the indicated siRNAs before being exposed to 1% O_2_ for 24h. **(E)** Quantitative RT–PCR analysis of CAIX mRNA prepared from HEK293 cells transfected with the indicated plasmids before being exposed to 1% O_2_ for 24h. (All Relative expression compared to RPL13A mRNA).

We next examines the potential for UCHL1 as a regulator of a well-described endogenous hypoxia-responsive gene CAIX[19]. CAIX transcript was found to be increased in cells exposed to low oxygen, as compared to control and this effect was attenuated by siRNA-mediated depletion of HIF1α, this consistent with CAIX being a HIF1-dependent target gene (Figure 2D). In cells transfected with UCHL1 and exposed to hypoxia, CAIX transcripts were increased as compared to control cells exposed to hypoxia alone (Figure 2E). These data are consistent with UCHL1 increasing the expression HIF-dependent genes through control of HIF1α stability.

### UCHL1-dependent control of HIF activity during liver fibrosis

The expression of UCHL1 is largely confined to brain and reproductive tissue in healthy individuals, however aberrant expression of UCHL1 can be pathogenic in various disease states [24, 25]. An example of this occurs during liver injury where UCHL1 transcript and protein expression is highly elevated during HSC transdifferentiation [18] (Figure 3A and 3B). As UCHL1 can deubiquitinate and stabilise HIF1α in cancer cell lines (needs a ref), we sought to determine if there are changes in HIF1α levels during HSC transdifferentiation. HIF1α levels were analysed from protein extracts prepared from cultures of primary human HSCs, a well-defined *in vitro* model for interrogating molecular events that occur during HSC transdifferentiation. Early (Day 1) cultured cells are representative of the qHSC, while later cultured cells (Day 10) can be considered as representative of the fibrogenic myofibroblast-like aHSC [9]. Immunoblot analysis revealed elevated HIF1α levels in activated HSCs, despite these experiments being performed at 21% O_2_ (Figure 3B). To examine if the observed increase in normoxic HIF1α levels in activated HSCs alters HIF activity we examined a panel of well-characterized HIF1α genes. cDNAs prepared from RNA isolated from human qHSC and aHSC were used to quantitate the transcript levels of the hypoxia-responsive genes; ANKRD37, CAIX, PHD2, PHD3, FGF and VEGF [19]. Each of the HIF targets was elevated in the activated HSCs consistent with the elevated levels of HIF1α (Figure 3C).

**Figure 3.**
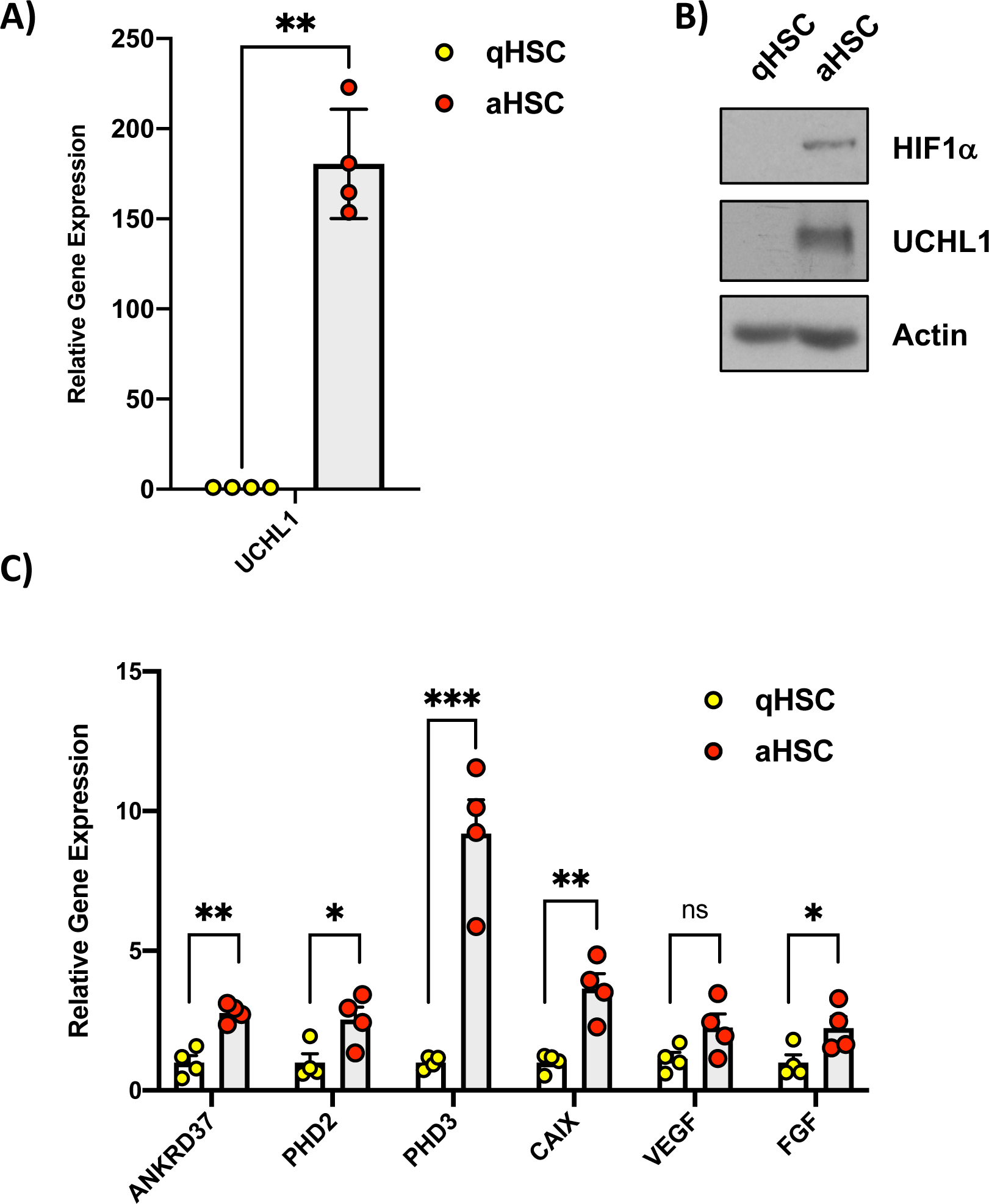
Expression of HIF1α and elevated HIF1 target gene expression in activated human hepatic stellate cells (HSCs). **(A)** Quantitative RT-PCR analysis of UCHL1 mRNA levels in from quiescent (Day 1) and activated (Day 10) HSCs All Relative expression compared to actin mRNA). **(B)** Immunoblot analysis of lysates prepared from quiescent (Day 1) and activated (Day 10) HSCs. **(C)** Quantitative RT-PCR of HIF1-target gene expression in quiescent (Day 1) an activated (Day 10) HSCs (All Relative expression compared to RPL13A mRNA).

### Genetic deletion or chemical inhibition of UCHL1 reduces HIF1α levels in hepatic stellate cells

To further investigate the relationship between UCHL1 levels and HIF activity we isolated protein extracts from HSCs derived from wildtype and UCHL1-deficient mice. Consistent with our previous observations, UCHL1 deletion was associated with reduced levels of HIF1α protein expression under normoxic conditions (Figure 4A). UCHL1-dependent control of HIF1α expression in *uchl1^-/-^* cells is post-transcriptional as WT and *uchl1^-/-^* HSCs express similar levels of HIF1α mRNA (Figure 4B). To investigate if reduced levels of HIF1α in UCHL1 deficient HSCs persists under hypoxic conditions, cells were incubated at 1% O_2_ for various time points. In wildtype HSCs there was an expected rapid and robust stabilization of HIF1α in response to hypoxic stress, however stabilization of HIF1α was muted in UCHL1 deficient cells (Figure 4C).

**Figure 4.**
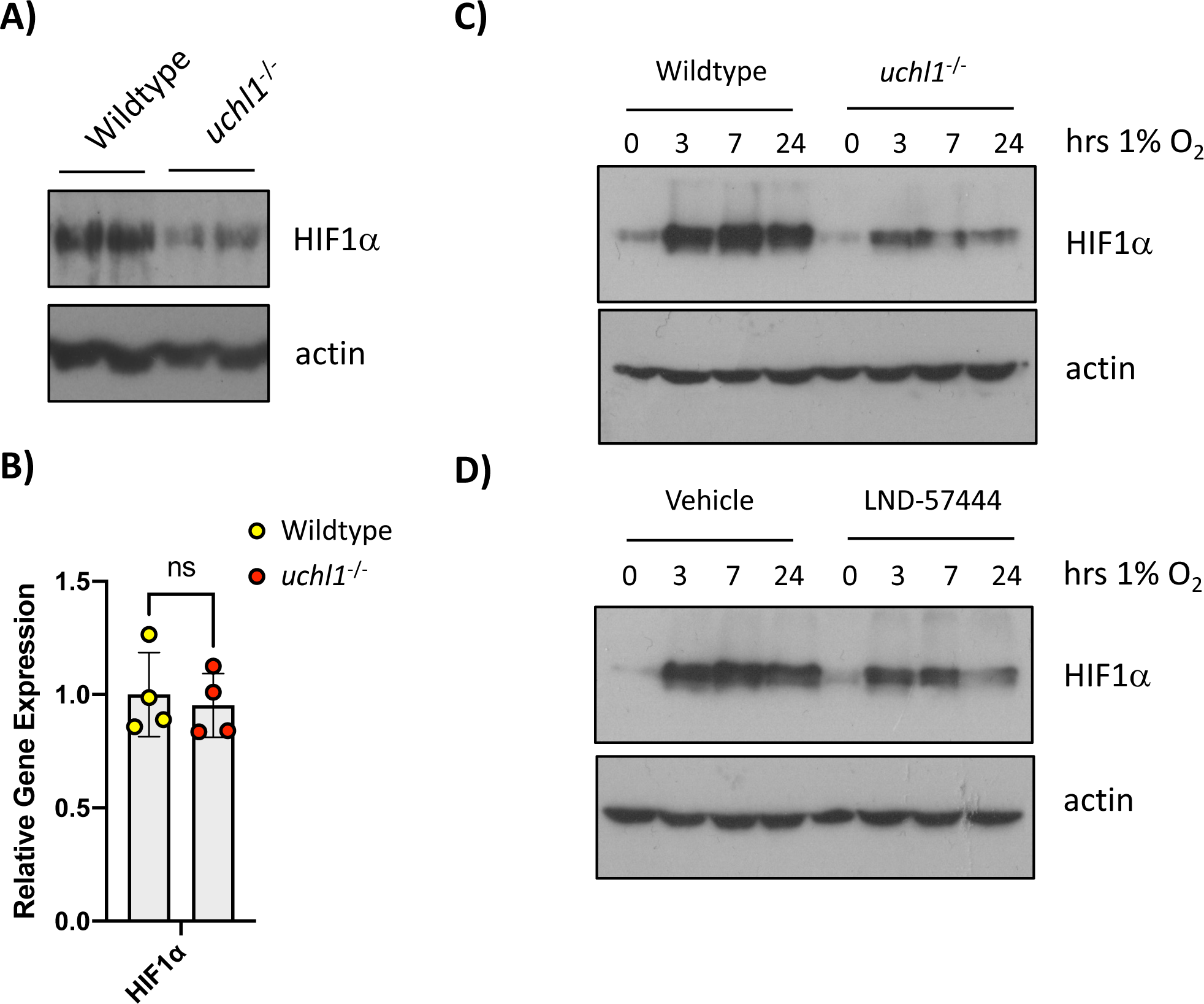
Reduction of HIF1α accumulation in UCHL1^-/-^ Cells. **(A)** Immunoblot analysis of lysates prepared from HSCs isolated from matched littermate wildtype and uchl1^-/-^ mice. **(B)** Quantitative RT-PCR analysis of Hif1a mRNA levels in from HSCs isolated from matched littermate wildtype and uchl1^-/-^ mice. (All Relative expression compared to RPL13A mRNA). **(C)** HSCs isolated from matched littermate wildtype and uchl1^-/-^ mice exposed to 1% O_2_ for the indicated times. Whole-cell lysates (WCLs) prepared from these cells were subjected to immunoblot analysis to assess expression levels of the indicated proteins. **(D)** HSCs prepared from wildtype mice were pre-treated with LDN 57444 (uM) for 30min before being exposed to 1% O_2_ for the indicated times. Whole-cell lysates (WCLs) prepared from these cells were subjected to immunoblot analysis to assess expression levels of the indicated proteins.

As germline deletion of any protein has the potential to lead to unanticipated changes and adaptation, we studied the effects of small molecule UCHL1 DUB inhibitors on HIF1α levels. To this end we used LDN-57444 a reversible, competitive, active-site directed inhibitor of UCHL1 [26]. Murine aHSCs were pre-treated with LDN-57444 and exposed to hypoxia for the indicated times. Pre-treatment with the UCHL1 inhibitor attenuated HIF1α stabilization by low oxygen, consistent with UCHL1 preventing ubiquitin-mediated degradation of HIF1α (Figure 4D).

### UCHL1 enhances the expression of HIF-responsive pro-fibrogenic gene targets

To investigate the relationship between HIF activity on pro-fibrotic gene expression in a tractable cell model, RNA was isolated from normoxic or hypoxic HEK293 cells and used to quantitate the transcript levels of COL1A1, SMA1, TGF*β*, and TIMP1, these transcripts representing well-defined markers of fibrgenesis. Levels of SMA1 and TIMP1 transcripts were increased in hypoxic cells, however no differences were seen for COL1A1 and TGF*β* mRNAs (Figure 5A). Depletion of HIF1α using specific RNAi prevented hypoxia-dependent increases in SMA1 and TIMP1 suggesting that HIF1 activity is required for these hypoxia-associated changes in gene expression (Figure 5A).

**Figure 5.**
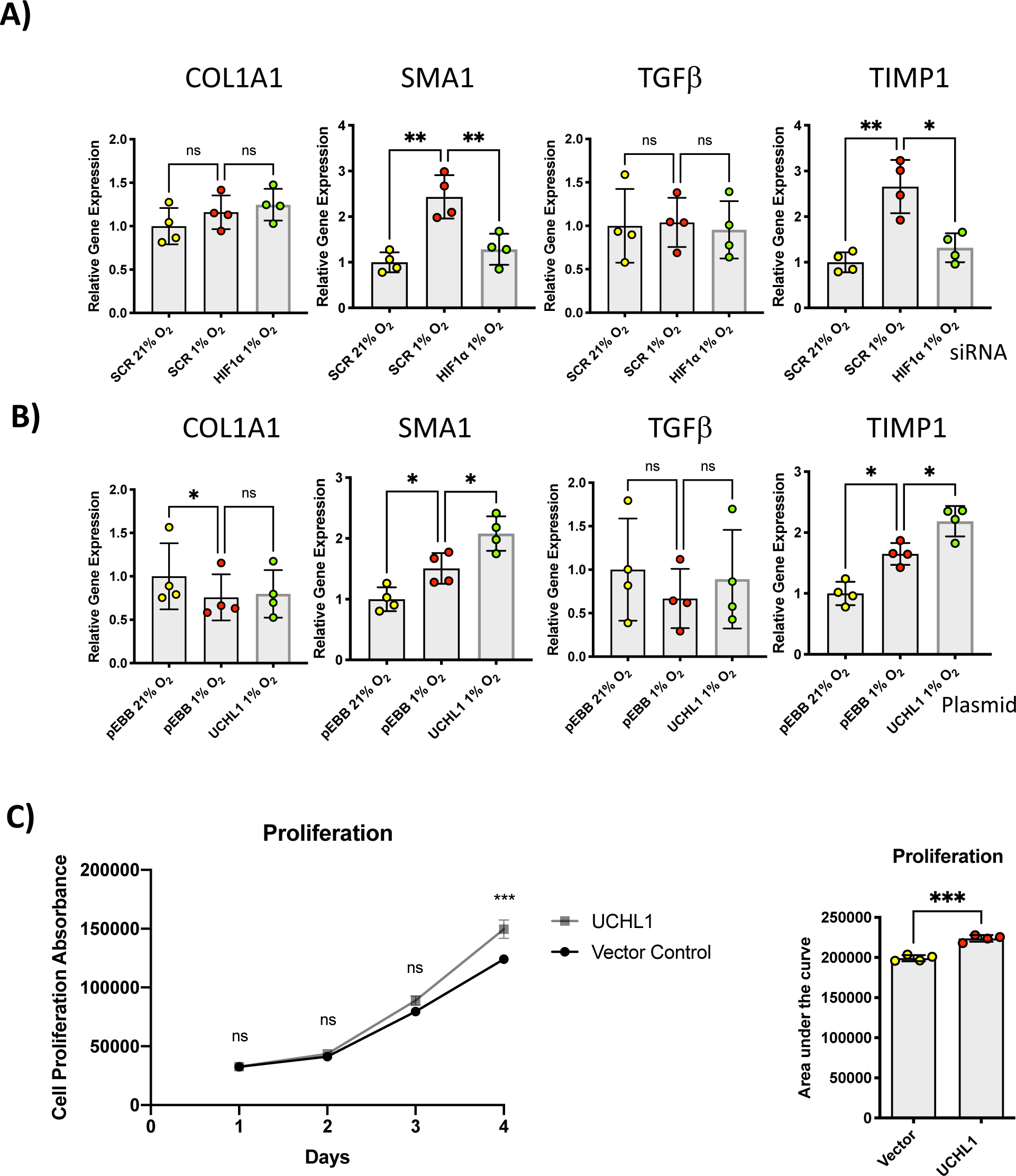
UCHL1 promotes the expression of HIF-responsive pro-fibrogenic gene targets. **(A)** Quantitative RT–PCR analysis of COL1A1, SMA1, TGFb and TIMP1 mRNA prepared from HEK293 cells transfected with the indicated siRNAs before being exposed to 1% O2 for 24h. **(B)** Quantitative RT–PCR analysis of COLA1, SMA1, TGFb and TIMP1 mRNA prepared from HEK293 cells transfected with the indicated plasmids before being exposed to 1% O2 for 24h. (All Relative expression compared to RPL13A mRNA). **(C)** Proliferation of HEK293 cells transfected with empty vector or UCHL1 as measured by PrestoBlue. Graph displays Raw absorbance values. **(D)** The area under the curve Raw absorbance x time(days)) was calculated and analysed by unpaired students t-test.

We next examined the relationship between UCHL1 and hypoxia-induced expression of pro-fibrotic genes. HEK293 cells were transiently transfected with UCHL1 and exposed to hypoxia for 24 hours prior to RNA isolation. Consistent with published reports expression of HIF-target gene expression was significantly induced by UCHL1 expression [15] (Figure 5B). Interestingly, only the HIF/hypoxia-responsive pro-fibrotic genes, SMA1 and TIMP1, were induced by UCHL1 overexpression consistent with the model of UCHL1 increasing HIF1α levels and HIF activity (Figure 5B).

To investigate the effect UCHL1 on cellular growth, cell proliferation rates were compared between HEK293 cells expression UCHL1 or transfected with vector alone (Figure 5C & 5D). HEK293 cells transfected with UCHL1 had a modest, but significant increase in growth rates, consistent with a role for UCHL1 as a regulator of cell proliferation (Figure 5C and 5D).

### UCHL1 Inhibition Reduces Fibrosis in Precision Cut Liver Slices

We next asked if inhibition of UCL1 effects fibrogenesis in a more physiological model of human liver fibrosis. To this end we selected hPCLS which involves culturing intact slices of human liver tissue in a rocked bioreactor which maintains liver structure, cellular composition and metabolic activity for up to 6 days, this offering a significant improvement on simple cell culture models.[21]. Fibrosis was induced in hPCLSs by addition of transforming growth factor beta 1 (TGFβ1) and platelet-derived growth factor (PDGFβ) to hPCLS media. UCHL1 activity in hPCLS was blocked under these conditions using two different concentrations (25 and 50μM) of LDN-57444. The higher concetration of LDN-57444 significantly inhibited TGFβ1+ PDGFβ stimulated increases in Collagen I and *α*SMA transcripts (Figure 6A and B respectively). Histological analysis of fibrogenesis in hPCLS was examined by *α*SMA and picosirus red staining (PRS) which detects cross-linking collagens. Exposure of hPCLS to TGFβ1+ PDGFβ increased *α*SMA and PRS staining with clear evidence of sinusoidal fibrosis in the latter stain, this strongly implicating *α*SMA+ aHSC as the major fibrogenic cells in this model. Remarkably, both concentrations of LDN-57444 prevented the accumulation of *α*SMA+ fibrogenic cells and deposition of cross-linking collagens (>Figure 6C). These effects were quantified for measurements made in hPSLS on 3 separate doner livers where in each case pharmacological inhibition of UCHL1 was associated with reduced fibrogenesis.

**Figure 6.**
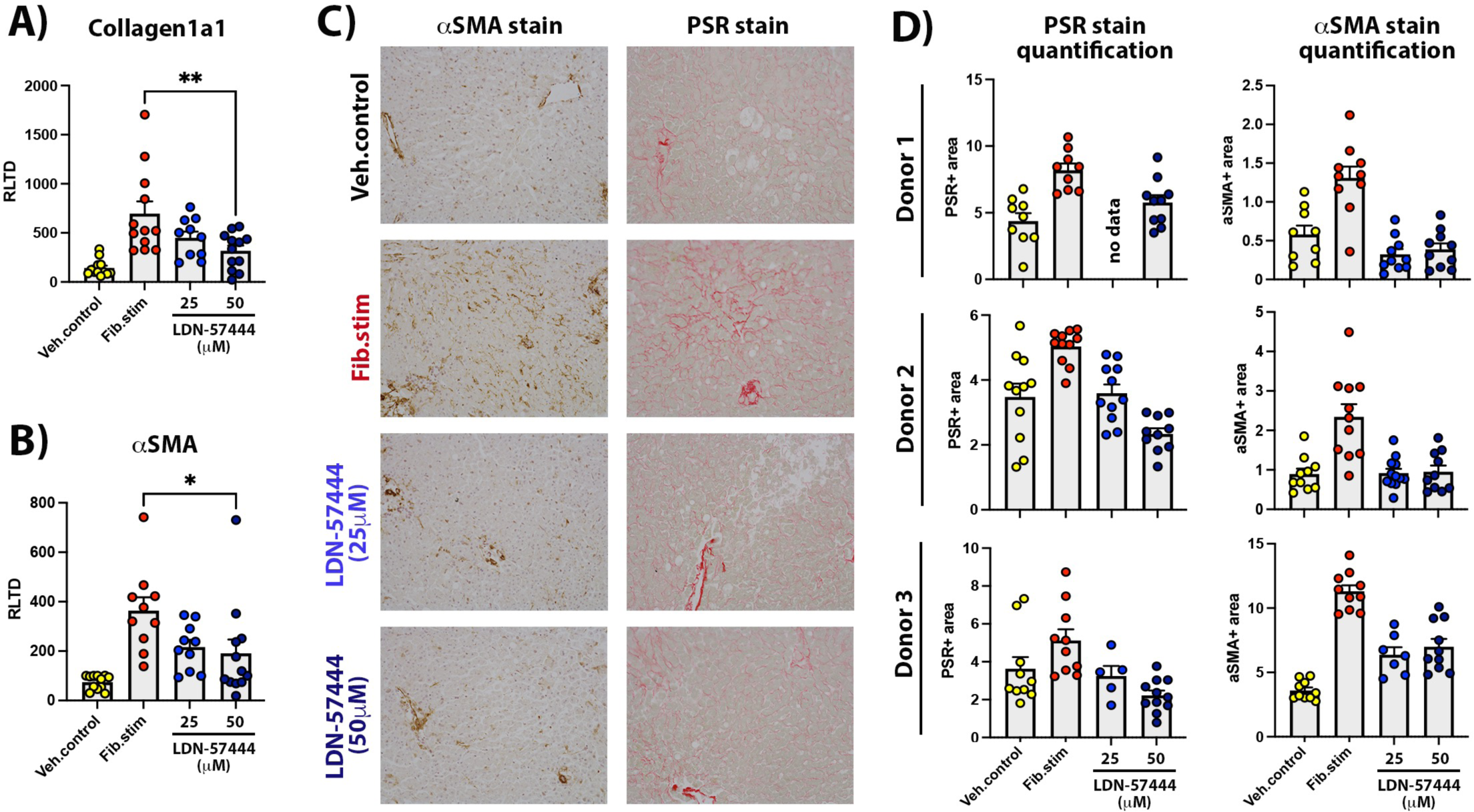
UCHL1 inhibitors have antifibrotic therapy in human liver slices. **(A)** mRNA levels of Col1A1 and αSMA in human PCLSs at t = 72h culture ± fib stim (TGFβ1/PDGFββ) ± LDN-57444 (96-hour total culture). Data are mean ± SEM in n = 3 different donor livers. *P < 0.05; **P < 0.01; ***P < 0.001; ****P < 0.0001. (B) Images of αSMA and picrosirius red–stained human PCLSs from representative donor liver at t = 72h culture ± fib stim (TGFβ1/PDGFββ) ± LDN-57444 (96-hour total culture). (C) Graphs show percentage area of picrosirius red–stained or αSMA stained tissue in human PCLS of individual donors at t = 72-hour culture ± fib stim (TGFβ1/PDGFββ) ± LDN-57444 (96-hour total culture). Abbreviations: Cont, control; fib stim, fibrotic stimulation with TGFβ1/PDGFββ; Veh, vehicle.

Our data collectively reveal that HIF1α expression and HIF1 activity are elevated in activated HSCs, even in conditions with adequate oxygen tensions. The high levels of UCHL1 expressed in activated HSCs specifically remove the degradative K48-linked ubiquitin chains from HIF1a, resulting in detectable HIF1α, even under normal oxygen tensions. The elevated HIF activity seen in HSCs results in elevated expression of pro-fibrogeneic HIF-target gene expression in activated HSCs.

## Discussion

A role for HIF1α in the pathogenesis of fibrosis has been well documented, with scarred, hypoxic tissue evident in diseased liver [27]. However, increases in HIF levels and activity often precede acute hypoxic stress, therefore alternative modes of HIF activation may exist to activate HIF in fibrotic tissue [27]. In this present study we describe a role for the deubiquitinating enzyme UCHL1 in controlling HIF activity in HSC under both normoxic and hypoxic conditions. Levels of UCHL1 were markedly elevated during HSC activation and this was associated with increased stability of HIF1α, and increased expression of HIF target genes (Figure 5). Genetic deletion or chemical inhibition of UCHL1 reduced HIF1α levels in activated HSCs under both normoxic and hypoxic conditions therby strongly implicating UCHL1 as an important regulator of HIF expression and activity.

HIF activity is mainly regulated by the sequential hydroxylation and ubiquitination of HIF1α and HIF2α by PHDs and VHL respectively, however there are a number of E3 ubiquitin ligases and DUBs that can alter HIF stability, activity and sub-cellular localisation [22]. UCHL1 is one such DUB that has recently been shown to promote HIF1α deubiquitination in a variety of cancer cell types [15, 17]. UCHL1 is highly expressed in the brain, where it is involved in cell death, learning, and memory (needs a ref). Dysregulation of has been linked to neurodegeneration [24], cancer [25, 28], and fibrosis-associated diseases [18, 29, 30]. Using small molecule inhibitors of UCHL1 we reveal that HIF signalling can be supressed, this potentially offering a therapeutic strategy to target HIF signalling in fibrotic tissue. In support of this idea, active fibrogenesis in a human pre-clinical model of human liver fibrosis was suppressed by UCHL1 inhibition (Figure 6). In this model, exposure of liver slices to the well-characterised autocrine and stimulators of fibrosis, TGFβ1+ PDGFβ, robustly induces fibrogenesis. Hence, demonstrating that pharamacological inhibtion of UCHL1/HIF signaling in this model is highly relevant for translation to human liver disease. Recent work using an alternative and more selective inhibitor of UCHL1, IMP-1710, revealed a reduction of expression of profibrogenic markers in fibroblasts isolated from patients with idiopathic pulmonary fibrosis, demonstrating the importance of UCHL1 signalling in fibrosis [31].

In addition to the link between UCHL1 and HIF in promoting the expression of pro-fibrogenic genes there is a clear link between UCHL1 controlling HIF activity in a variety of tumour types [15–17]. UCHL1 activity correlates with the poor prognosis of patients with breast and lung cancers, and a strong correlation between UCHL1 levels and HIF activity were observed in patient samples[15]. It would be of great value to investigate the efficacy of UCHL1 inhibitors in treating cancer cell types with elevated UCHL1 and HIF activity.

## Abbreviations

HSC: Hepatic Stellate Cell
DUB: Deubiquitinase
HIF: hypoxia inducible factor
UCHL1: Ubiquitin C-terminal hydrolase-L1.

## Acknowledgements

Funding for this work was provided by Newcastle University Independent Research Establishment Scheme Fellowship to NSK, North West Cancer Research Development Grant [RDG2021.12 to NSK]. BBSRC DTP Studentship [BB/T008695/1 to GBE].

## Competing Interests

DM is a director of Fibrofind. JL and DM are shareholders in Fibrofind limited.

